# Real-time prediction of drug-induced proarrhythmic risk with sex-specific cardiac emulators

**DOI:** 10.1101/2024.09.30.615798

**Authors:** Paula Dominguez-Gomez, Alberto Zingaro, Laura Baldo-Canut, Caterina Balzotti, Borje Darpo, Christopher Morton, Mariano Vázquez, Jazmin Aguado-Sierra

## Abstract

*In silico* trials for drug safety assessment require a large number of high-fidelity 3D cardiac electrophysiological simulations to predict drug-induced QT interval prolongation, making the process computationally expensive and time-consuming. These simulations, while necessary to accurately model the complex physiological conditions of the human heart, are often cost-prohibitive when scaled to large populations or diverse conditions. To overcome this challenge, we develop sex-specific emulators for the real-time prediction of QT interval prolongation, with separate models for each sex. Building an extensive dataset from 900 simulations allows us to show the superior sensitivity of 3D models over 0D single-cell models in detecting abnormal electrical propagation in response to drug effects as the risk level increases. The resulting emulators trained on this dataset showed high accuracy level, with an average relative error of 4% compared to simulation results. This enables global sensitivity analysis and the replication of *in silico* cardiac safety clinical trials with accuracy comparable to that of simulations when validated against *in vivo* data. With our emulators, we carry out *in silico* clinical trials in seconds on a standard laptop, drastically reducing computational time compared to traditional high-performance computing methods. This efficiency enables the rapid testing of drugs across multiple concentration ranges without additional computational cost. This approach directly addresses several key challenges faced by the biopharmaceutical industry: optimizing trial designs, accounting for variability in biological assays, and enabling rapid, cost-effective drug safety evaluations. By integrating these emulators into the drug development process, we can enhance the reliability of preclinical assessments, streamline regulatory submissions, and advance the practical application of digital twins in biomedicine. This work represents a significant step toward more efficient and personalized drug development, ultimately benefiting patient safety and accelerating the path to market.

## 1 Introduction

Arrhythmias manifest as irregular heart rhythms stemming from abnormal cardiac electrical activity, ranging from benign palpitations to severe conditions like ventricular fibrillation [1]. A pivotal biomarker associated with increased arrhythmic risk is a prolonged QT interval, as measured on electrocardiograms (ECGs) from the beginning of the QRS complex to the end of the T wave. The change from baseline QT, the ΔQT, is notably linked to torsades de pointes (TdP), a potentially lethal, polymorphic ventricular tachycardia [2]. Prolongation of the QT interval, caused by many drugs beyond cardiac treatments, often arises from unintended ion channel blockades such as the hERG potassium channel, crucial for cardiac repolarization. Such blockade delays repolarization, thereby prolonging the QT interval. The gold standard for assessing the proarrhythmic risk of a drug traditionally involves measuring its effects on the hERG channel *in vitro*, a single-dose safety pharmacology study in dogs or monkeys, and evaluation of potential effects on the QT interval in clinical trials. Despite substantial research on this topic [3–6], the precise relationship between drug-induced ion channel blockade at the cellular level and QT prolongation at the organ level remains poorly understood, complicating risk assessments.

Regulatory agencies, such as the US Food and Drug Administration (FDA) and the European Medicines Agency, have recognized the potential of computational models and digital twins to enhance drug safety evaluations, as they offer a promising approach to bridge the gaps in the translation of nonclinical assays to clinical outcomes in the drug discovery phase and in the early preclinical stage [6]. The Comprehensive in Vitro Proarrhythmia Assay (CiPA) initiative, for instance, promotes the use of *in silico* models alongside *in vitro* and clinical data to complement proarrhythmic risk assessment [7].

Computational models for predicting proarrhythmic risk vary in complexity [8]. At the simplest level, single-cell 0D models (zero dimensional, where the spatial dependence of variables is neglected in favor of time dependence only) simulate the electrical activity of individual cardiac cells, providing insights into ion channel behavior and drug effects. However, these 0D models cannot capture QT interval prolongation as they do not account for the holistic behavior of the heart. Given that QT interval measurement is the gold standard for assessing proarrhythmic risk in clinical practice, employing 3D cardiac models for *in silico* clinical trials is essential. These models account for the intricate anatomy of the heart and simulate cardiac function at the organ level, providing a more comprehensive and accurate representation of drug-induced effects on cardiac electrophysiology. Several studies have predicted proarrhythmic risk using 0D models [9–14], while others have advanced to 3D representations of the heart for assessing ΔQT [15–21]. However, gaps persist in the literature in systematically determining the necessity of 3D models for accurate drug-induced proarrhythmic risk prediction compared to 0D single-cell models.

In addition, despite several advancements in 3D cardiac models, very few models account for sex-specific differences [21–24], even though substantial evidence indicates that female are more susceptible to arrhythmias [25]. While incorporating sex differences may be less critical for drugs causing only mild QT prolongation at therapeutic doses, it becomes essential for evaluating drugs with a high risk of TdP or substantially differing absorption rates between sexes. This gap underscores the urgent need to integrate anatomical and phenotypical sex differences into proarrhythmic risk predictions.

Despite their high accuracy, detailed cardiac 3D models can pose significant computational challenges for application in large-scale or real-time contexts. For drug discovery and at the preclinical level, the ability to predict ΔQT in real-time is crucial for gaining an early understanding of a drug’s safety profile, identifying safety margins, and streamlining subsequent development stages. This capability facilitates protocol modifications, dose regimen adjustments, or prompt discontinuation of problematic drugs, thereby optimizing resource allocation and ensuring that only the most promising drug candidates move forward. Advancing the concept of digital twins in biomedicine requires accelerating numerical simulations to achieve real-time virtual representations of living systems [26]. In the context of clinical trials, such advancements could enable rapid, real-time screening methods, particularly during the preclinical stage. These digital twins could act as instant-response systems for designing computational trials, allowing for faster and more efficient evaluations of drug safety and efficacy.

To address these challenges, surrogate models based on artificial intelligence methods, offer a promising solution. These surrogate models, or emulators, approximate the outputs of high-fidelity simulations at a fraction of the computational cost. Previous works have demonstrated the utility of emulators in cardiac modeling [27–35] and drug safety assessment [36, 37]. Notably, Costabal *et al*. [36] have pioneered the development of an emulator for ΔQT prediction, trained on a combination of 3D and 1D electrophysiological simulations.

In this study, we build on our previous work [23] by developing two emulators for real-time proarrhythmic risk assessment, specifically for ΔQT estimation. Each emulator is tailored for each sex, and is trained exclusively using data from 3D simulations. This process involved 900 electrophysiological runs, requiring approximately 2.1 million CPU hours globally. Prior to developing the emulators, we compared results from the 3D and 0D models. Our analysis demonstrated that 3D modeling is significantly more sensitive than 0D modeling in capturing the onset of abnormal electrical propagation in the context of proarrhythmic risk assessment for drugs, especially as the risk level increases. Leveraging these insights, we developed the emulators which achieved high accuracy with an average relative error of less than 4% compared to the simulator results, and a computational speed-up of five orders of magnitude. We then applied the emulators to replicate clinical trials for four benchmark drugs, comparing their predictions with both simulator results from different anatomies and real clinical data. This comparison validated the accuracy and generalizability of the emulators, demonstrating also their reliability in practical scenarios.

## 2 Results

We present a schematic overview of the general methodology used to build our emulators for realtime cardiac ΔQT prediction (see Figure 1). The emulators are generated using data derived from 3D high-fidelity electrophysiological simulations (i.e., obtained with the simulator). To simulate the drugs’ effect, we sample blockades of the seven most relevant ionic channels for cardiac proarrhythmic assessment [38]: *I*_CaL_, *I*_NaL_, *I*_to_, *I*_Ks_, *I*_K1_, *I*_Na_, and *I*_Kr_. We account for sex differences by employing detailed biventricular geometries for both male and female patients, and include respective phenotypes [23]. Using Alya, our finite element simulator [39], we performed 3D simulations to compute the ECG and to measure QT interval prolongation relative to a baseline configuration (ΔQT). This data is then used to train our emulators. We develop two separate emulators, one for males and one for females, each predicting ΔQT in real-time based on the blockades of the seven ionic channels. Each emulator comprises a classifier and a Gaussian process regression (GPR) model. The former classifies the data into non-arrhythmic or arrhythmic; the latter predicts ΔQT. With the term “arrhythmic”, in the context of these emulators, we refer to cases where the ΔQT cannot be computed due to an abnormal electrical propagation pattern in the ECG or where the ΔQT computed *in silico* exceeds a threshold [40]. The last criterion excludes cases producing values much larger than those typically measured in clinical practice and associated with the presence of arrhythmic events.

**Figure 1:**
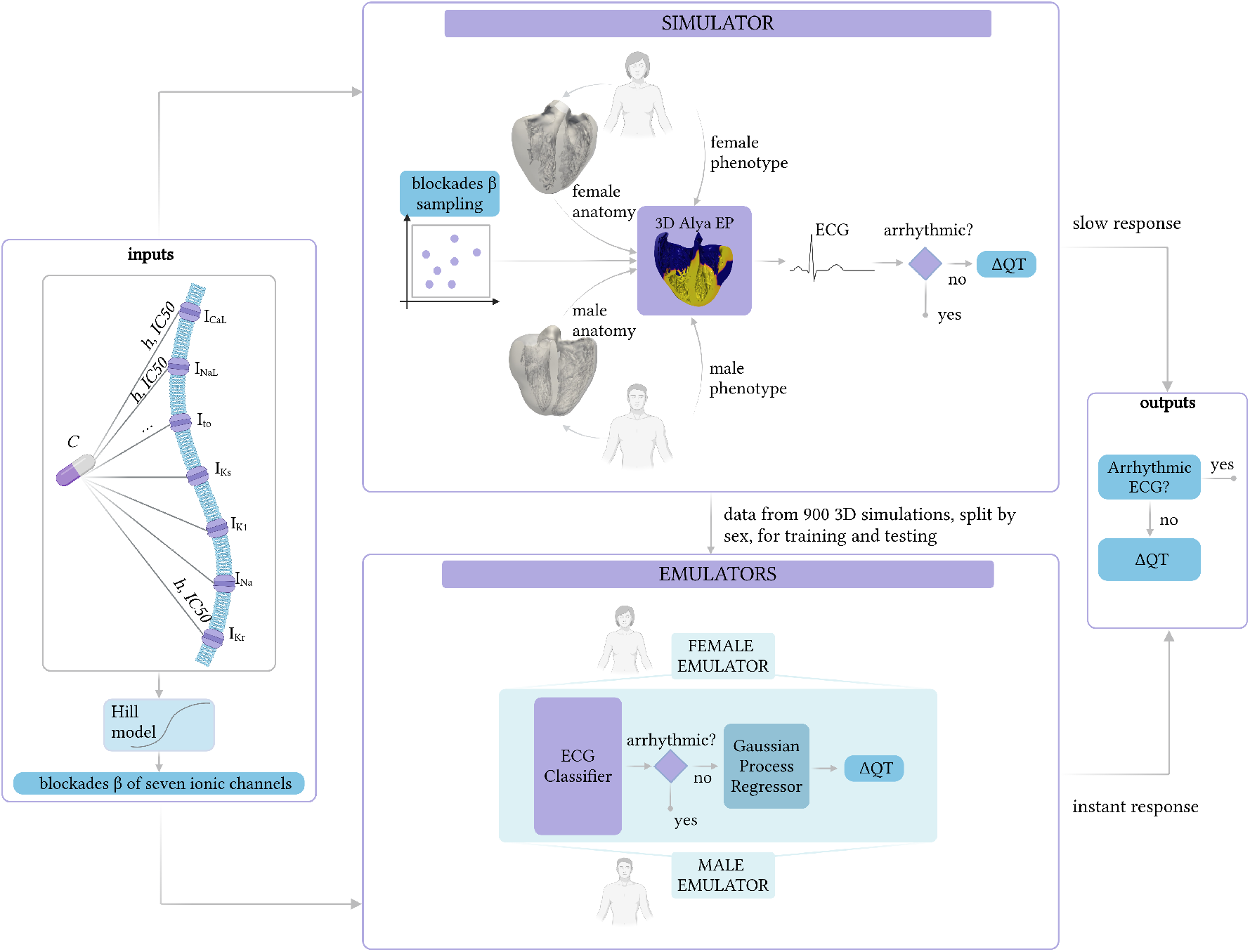
Overview of the methodology employed to build the ΔQT emulators.

### 2.1 Simulator results

In Figure 2, we present the simulation data that we obtain, and that we then use to train our emulators. This dataset comprises 900 simulations, with 450 simulations per sex, encompassing both arrhythmic and non-arrhythmic cases. To create our dataset, we excluded all the cases in which the 0D O’Hara-Rudy (ORd) initialization model [41] did not converge, i.e., it did not produce a periodic solution in terms of calcium concentration of each cell type (0 cases for males, 37 for females). Figure 2(a) displays the ECG signals, while Figure 2(b) presents the distributions resultant from the computation of ΔQT values for the entire dataset. Notably, the presence of arrhythmia precludes the computation of ΔQT; consequently, these cases are not included in the ΔQT distributions but their ECG patterns are depicted in grey in Figure 2(a). The results in this plot suggest that, on average, women tend to have longer QT interval durations. Additionally, Figure 2(b) shows a broader range of ΔQT values for men, likely because women are more prone to developing arrhythmic behavior under the same drug conditions. As a result, their data are more often excluded from the distribution. Figure 3 showcases examples for both males and females, highlighting differences through electrical propagation maps and corresponding ECG patterns based on the presence and absence of arrhythmias. It is worth noting that 11% of the female cohort exhibited arrhythmic behavior, compared to only 2% of the male cohort. The higher prevalence of non-convergent and arrhythmic cases among female subjects resulted in a reduced female sample size, accounting for the difference in distribution areas between males and females in Figure 2(b).

**Figure 2:**
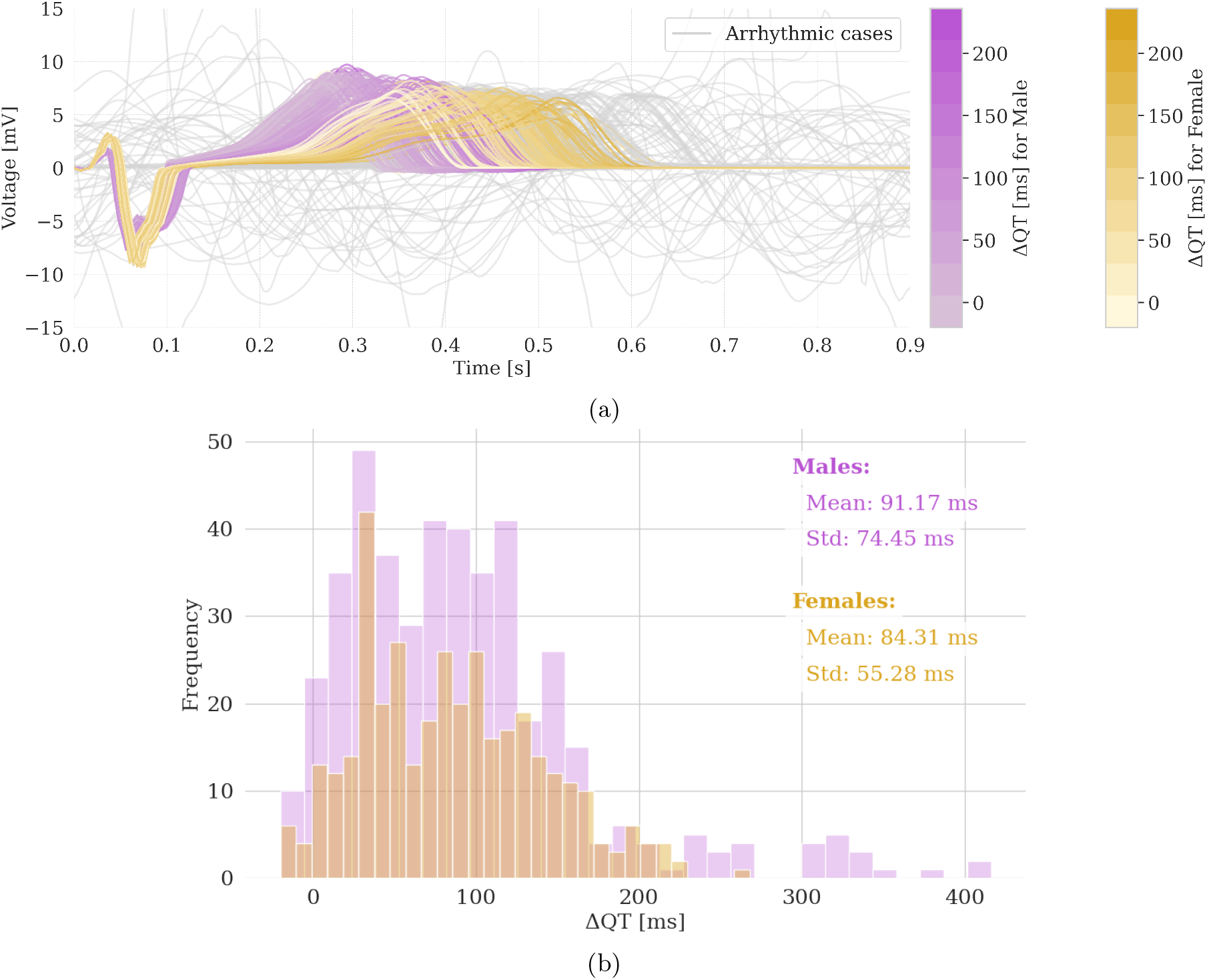
Simulator results. (a) ECG lead I signals for male and female subjects, with grey signals representing arrhythmic cases. (b) ΔQT distributions for male and female subjects.

**Figure 3:**
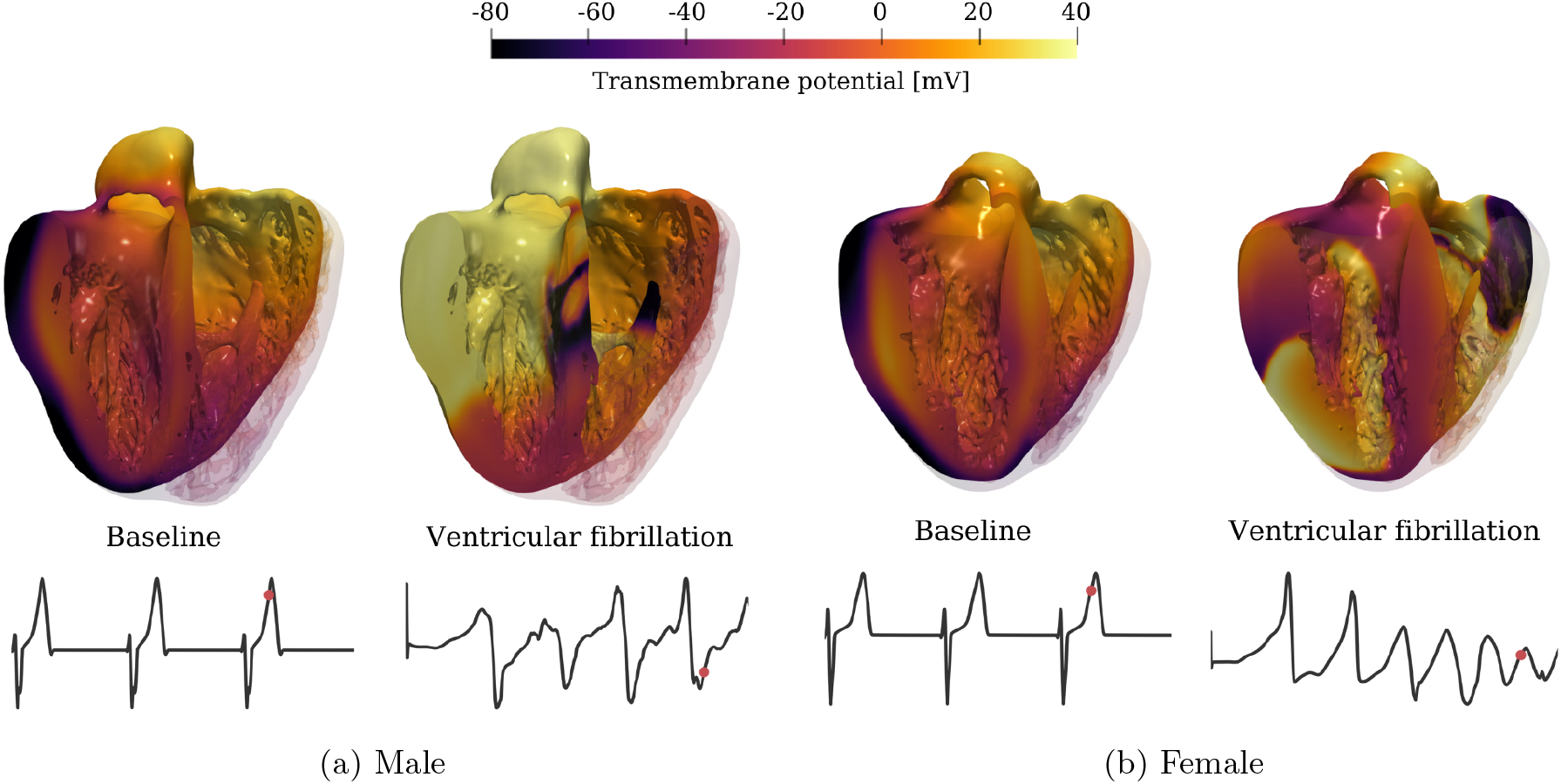
Examples of 3D electrophysiological simulations obtained with detailed biventricular anatomies. Both for the male (a) and female (b) anatomies, we report on the left the baseline simulation (without drug), and on the right the simulation in ventricular fibrillation conditions (due to drug). The signals represent the computed ECG lead I for each simulation and the red point on the ECG curve denotes the time in which the 3D images are taken.

### 2.2 Modeling proarrhythmic risk in 0D and 3D

3D cardiac models, while computationally expensive, capture spatial details and allow for the computation of ΔQT, aligning with clinical outcomes. In contrast, 0D models rely on metrics that cannot be measured clinically, such as the action potential duration (APD) or qNet (computational biomarker based on the sum of the net ionic currents over the course of the action potential [42]). To assess the value added by 3D models, we analyze the relationship between 3D and 0D simulation outcomes for evaluating proarrhythmic risk, as depicted in Figures 4 and 5. The 0D results are obtained from the initialization of the 3D simulation with the 0D ORd model run in endocardial, mid-myocardial, and epicardial cells.

**Figure 4:**
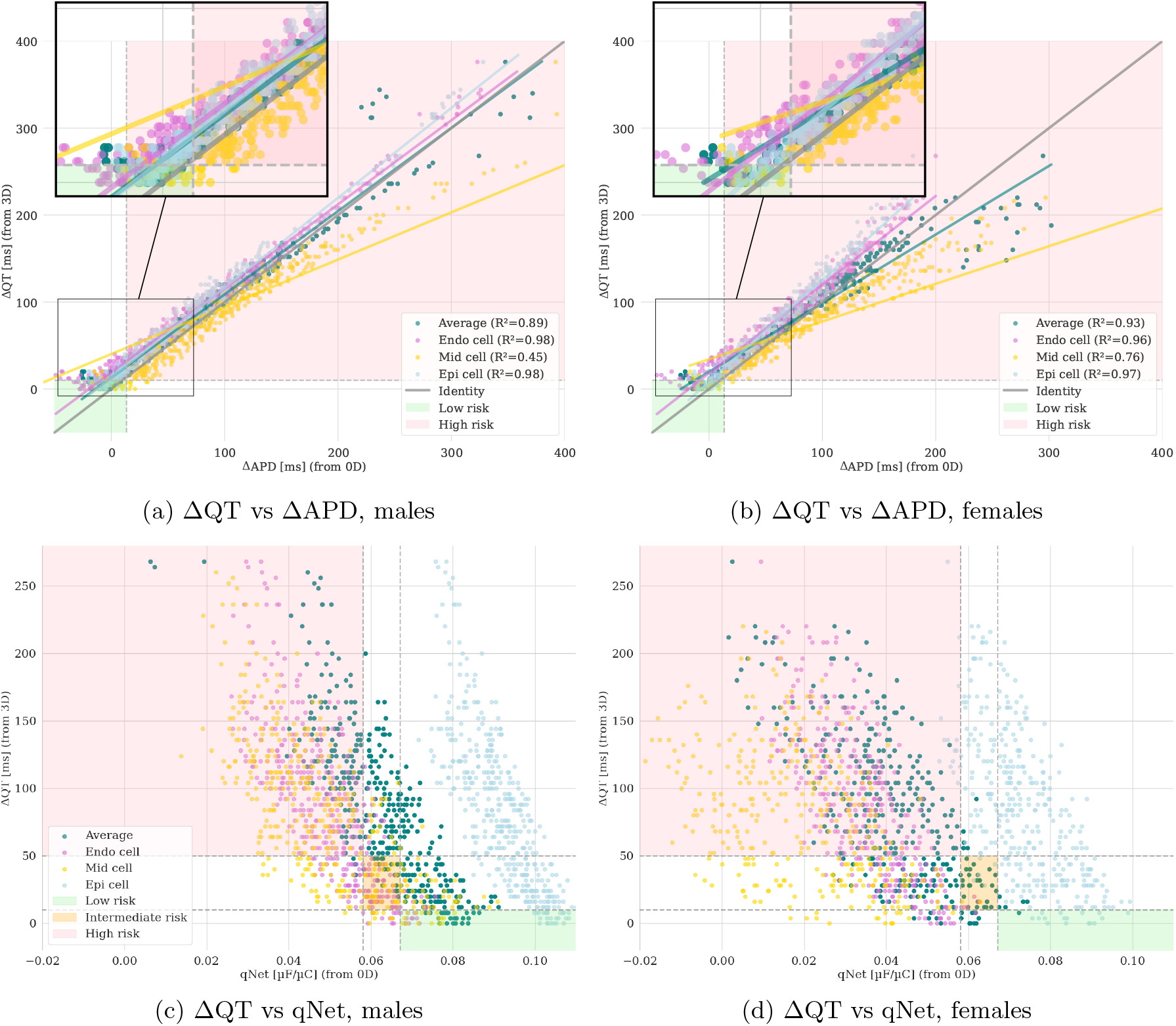
Comparison of biomarkers from 3D and 0D simulations to assess proarrhythmic risk. Data points are colored according to the cellular type: endocardial (Endo cell), mid-myocardial (Mid cell) and epicardial (Epi cell) types. Average represents the average of the three cellular types. Linear regression lines for each cell type indicate the general trend, and corresponding R^2^ scores are reported in the legends. The shaded areas mark the low, intermediate, and high-risk regions.

**Figure 5:**
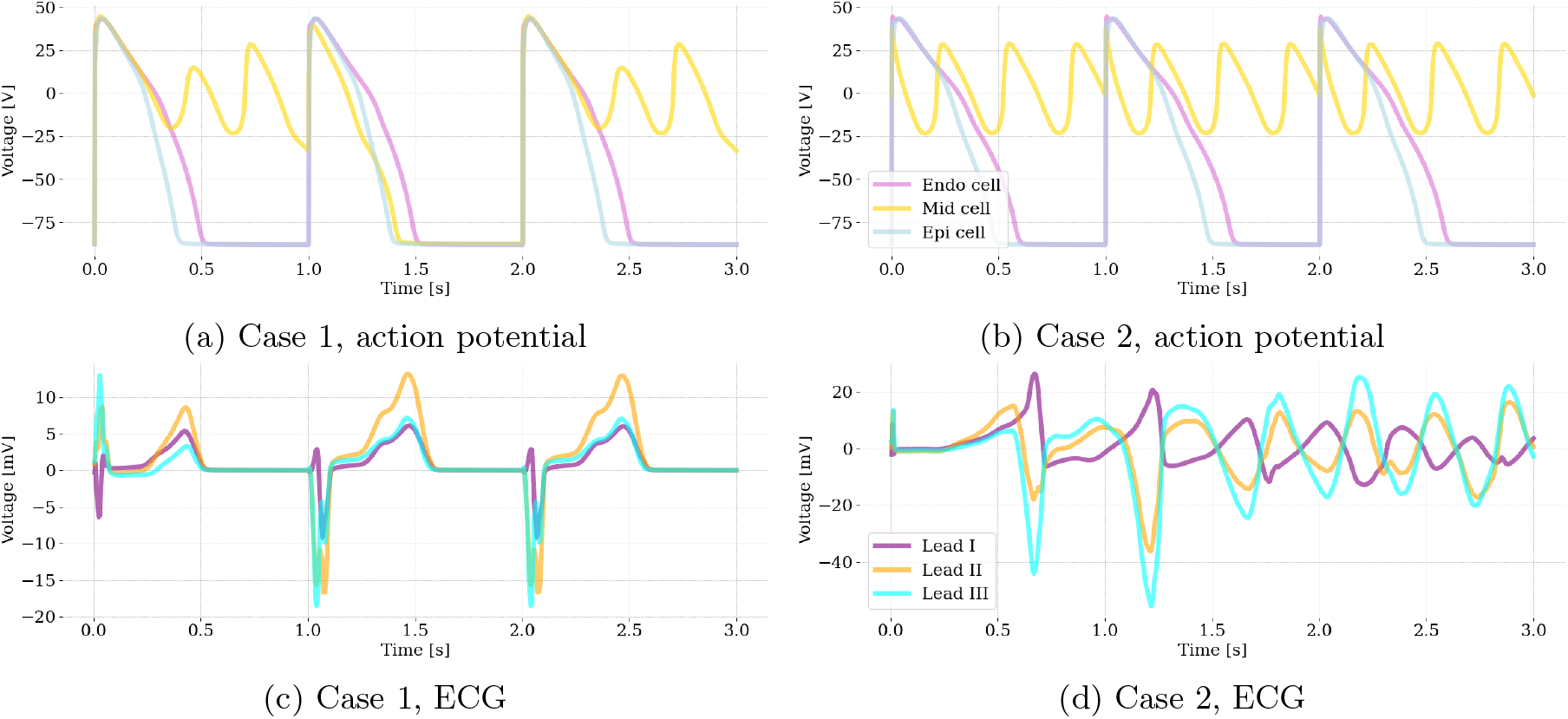
Comparison of two cases with similar action potentials leading to different ECG outcomes. Action potentials in (a) and (b) with abnormal behavior of the mid cell, are obtained from the last 3 beats of the 0D model (serving as initialization of the 3D simulation). However, the ECGs show different responses: (c) presents QT prolongation (170 ms) and ST elevation, while (d) manifests ventricular fibrillation. This illustrates that very similar cellular-level events can produce different effects on overall cardiac electrophysiological function.

Figure 4 illustrates a comparison of biomarkers from 3D and 0D simulations, with 4(a) and 4(b) depicting the relationship between ΔAPD (i.e., the difference between APD after and before drug administration, computed for different cell types) and ΔQT for females and males, respectively. The data points are differentiated according to the cell types: endocardial (Endo), mid-myocardial (Mid), and epicardial (Epi) cells. The colored areas represent low and high-risk regions. The highrisk threshold for ΔQT is set at 10 ms as it is commonly accepted that changes in QT interval of less than this value are generally considered to be within the normal physiological range [43], and for ΔAPD at 13.4 ms. This threshold is based on the established relationship where QT change is approximately 1.34 times the APD change, as reported by Mirams *et al*. [44]. Linear regression lines are fitted for each cell type, along with an overall average. The relationship between ΔAPD and ΔQT appears linear at lower values, but the data points spread as the values increase, indicating that the linear regression model’s fit decreases with higher risk. This trend is quantified by the R^2^ scores [45], which show the extent to which the model fits the data linearly. The R^2^ scores for the overall average and individual cell types highlight that while there is a general linear trend, significant deviations occur at higher risk levels. To further investigate this, we performed a cumulative sum test to assess the loss of linearity in the 0D model predictions compared to the 3D predictions. The test revealed that for ΔQT values of 100 ms in females and 125 ms in males, the residuals increase significantly and begin to diverge. This divergence suggests a significant difference in behavior between the 0D and 3D models beyond these thresholds, marking the onset of what we refer to as the arrhythmogenic window. The analysis suggests that while 0D models can reliably predict outcomes below this arrhythmogenic window, their ability to detect electrical propagation abnormalities diminishes considerably within the window as the risk level rises, when compared to the 3D model.

Figures 4(c) and 4(d) illustrate the relationship between qNet and ΔQT for females and males, respectively. qNet is intended to classify the risk of drug-induced arrhythmias into low, intermediate, and high-risk categories by evaluating the net charge carried by ions during the cardiac action potential. The plots delimit these three risk categories (intermediate risk: qNet ∈ [0.0581, 0.0671] μF*/*μC and ΔQT ∈ [10, 50] ms), by coloring the different corresponding regions. These thresholds are based on the FDA’s CiPA initiative guidelines [46] and statistics on drug-induced risk of lifethreatening arrhythmias [47]. Despite its utility for risk classification, qNet shows limited accuracy and fails to predict risk correctly for any female cell type. This constraint is evident in the plots, where the classification boundaries do not align well with the actual data points for females: our results suggest that qNet often misclassifies female cell types into incorrect risk categories.

Figure 5 compares two cases with similar action potentials (from the 0D model) leading to different ECG outcomes (from the 3D model). Figures 5(a) and 5(b) show the action potentials for two subjects with abnormal mid-cell behavior. Despite similar cellular-level events, the ECG signals in Figures 5(c) and 5(d) demonstrate significantly different outcomes: case 1 exhibits QT prolongation, while case 2 experiences ventricular fibrillation. These comparisons highlight that even with similar cellular behaviors, the overall cardiac outcomes can vary greatly. QT prolongation of case 1 suggests a significant but non-lethal proarrhythmic risk, whereas ventricular fibrillation characterizing case 2 indicates a life-threatening condition. The shortcomings of 0D models become particularly apparent under conditions of high QT prolongation, where they fail to capture the complexities of whole heart interactions and tissue-level electrical impulse propagation that are crucial for accurate risk assessment.

### 2.3 Accuracy of the emulators compared to simulator results

First, a preliminary version of the emulators is introduced and used for database design with a conservative approach to the ion channels’ blockade range. Then, a global sensitivity analysis (GSA) is performed to understand the main ion channels that influence our model response and consequently to expand and refine our database. This provides the ability to create an enhanced version of the emulators based on this last dataset, which are evaluated for their accuracy and computational performance in comparison to the simulator. With this, the emulators are capable of predicting the ΔQT responses even with higher blockades of the ion channels, promoting the occurrence of arrhythmic events.

#### 2.3.1 Emulator preliminary version

We begin by designing the dataset for predicting ΔQT using a GPR model. Denoting by *N* the sample size, the input of the GPR model are vectors ***x***_*i*_ ∈ ℝ^7^, *i* = 1, *…, N*, containing the curren blockades of the seven ionic channels. The output *y*_*i*_ ∈ ℝ, *i* = 1, *…, N*, is the corresponding ΔQT value.

We consider blockades of the seven channels *β*_k_ ∈ [0.0, 0.6], with k = CaL, NaL, to, Ks, K1, Na, Kr, as done in [36]. The input vectors are selected by performing Latin hypercube sampling [48] as *N* varies in *{*150, 250, 350*}*. To compare the results obtained for the different values of *N* and determine the ideal sample size, we fix the test set as 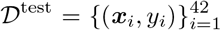 for all the three cases, where 42 represents the 28% of the data when *N* = 150. The training set 𝒟^train^ consists of 72% of the remaining part of the dataset 𝒟 *\* 𝒟^test^. Then, for each *N*, we tune the sex-specific optimal hyperparameters through an automatic exhaustive search that maximizes the R^2^ score. Table 1 presents the variation in the sample size *N* of the R^2^ score, the mean absolute error *ε*^MAE^, the mean absolute percentage error *ε*^MAPE^, and the root mean squared error *ε*^RMSE^. The scores are high for both sexes, regardless of *N*. In the male case, the errors show less sensitivity to *N*, while they exhibit a considerable improvement as *N* increases for females.

**Table 1:** R^2^ scores and prediction errors on the test set 𝒟^test^ for males (M) and females (F) using different values of *N*.

| $N$ | $R^2$ score | | $\epsilon^{\text{MAE}}$ [ms] | | $\epsilon^{\text{MAPE}}$ [%] | | $\epsilon^{\text{RMSE}}$ [ms] | |
| --- | --- | --- | --- | --- | --- | --- | --- | --- |
|  | M | F | M | F | M | F | M | F |
| 150 | 0.999 | 0.995 | 1.409 | 3.052 | 3.364 | 5.851 | 1.680 | 3.961 |
| 250 | 0.999 | 0.998 | 1.384 | 1.895 | 3.575 | 3.997 | 1.750 | 2.557 |
| 350 | 0.999 | 0.998 | 1.349 | 1.877 | 3.411 | 3.829 | 1.705 | 2.412 |

The GPR model, tuned with *N* = 350, represents the preliminary version of the emulators used to perform the GSA [49]. The objective is to identify the ionic channels that most significantly impact the output, specifically focusing on inputs that predominantly induce a high risk of QT prolongation. Based on [47] (as we also do in the analysis of Figure 4), we fix the high-risk threshold to 50 ms and use the GSA to examine the channels that primarily contribute to achieving a ΔQT exceeding this threshold. As shown in Table 2, the three most influential ionic channels for both males and females are *I*_Kr_, *I*_NaL_, and *I*_Ks_, with *I*_Kr_ providing the most substantial contribution, both individually and through interactions with other channels.

**Table 2:**
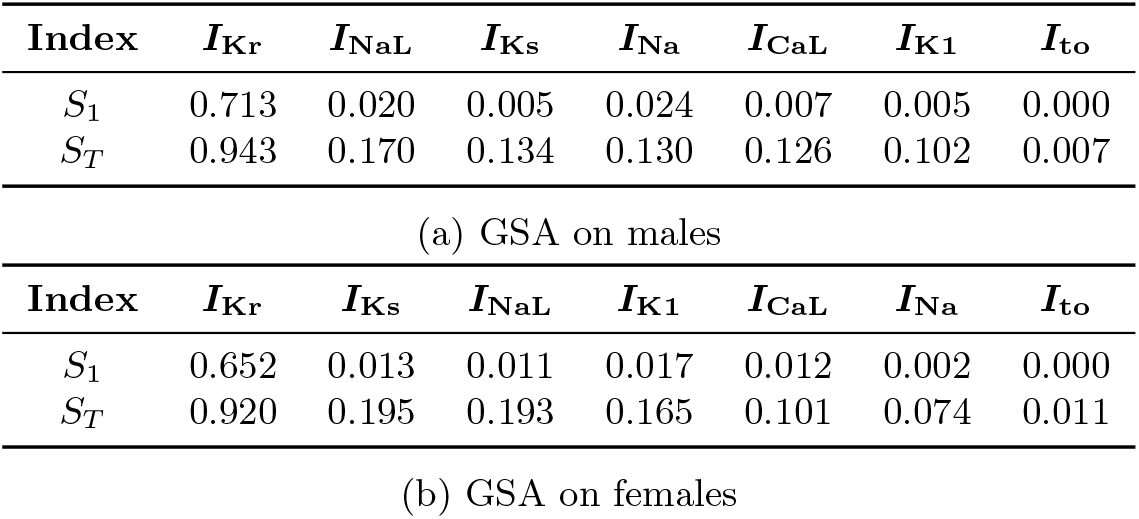
GSA with the preliminary version of the emulators (*β*_k_ ∈ [0.0, 0.6], with k = CaL, NaL, to, Ks, K1, Na, Kr). *S*_1_ and *S*_*T*_ are the first and total Sobol’ indices [49], respectively.

In light of this, we leverage the results of the GSA to expand our database. Specifically, we conduct an additional 100 simulations per sex, focusing on the blockades of these three critical ion channels *β*_k_ ∈ [0.4, 0.9], with k = Kr, NaL, Ks, while setting the remaining blockades to zero. Throughout the rest of the paper, we present results from these enhanced emulators.

#### 2.3.2 Emulator enhanced version

Expanding the dataset with higher blockade levels leads to an increase in arrhythmic cases. To predict ΔQT, we design our enhanced emulator using a classifier followed by a GPR model. The classifier enables us to filter out inputs that lead to arrhythmias so that we train the regressor only on relevant data.

##### ECG Classifier

The first step of the emulators consists of a classifier to determine whether the ECG is arrhythmic: it is characterized by an abnormal signal – for which the ΔQT is not computable – or the QT prolongation exceeds a fixed threshold *τ*. Specifically, following [40], we set *τ* = 240 ms and *τ* = 196 ms for males and females, respectively. This choice allows us to limit the QT to the threshold value of 600 ms for both sexes.

Considering the same inputs in Section 2.3.1, the output of the classifier is 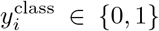, *i* = 1, *…, N*, where a value of 0 (negative) indicates a non-arrhythmic case, while 1 (positive) indicates an arrhythmic one. In our data, 92% of the outputs are negative for males, compared to 85% for females. We randomly divide our data into a training and test set, allocating 72% of data for training and 28% for testing. Note that the data is split in a way that preserves the proportion of positive and negative values in both the training and test sets. We use XGBoost Classifier (XGBC) [50] and *k*-Neighbors Classifier (KNC) [51] for males and females, respectively. We tune the optimal hyperparameters on the training set through an exhaustive search, looking for those that maximize the F1 score [52], as shown in Table 3. The performance of the classifier is reported in Table 4. The evaluation of accuracy, precision, recall, and F1 score metrics [52] on the test set reveals perfect scores for males and consistently high scores for females. The classifier’s strong performance is further illustrated in Figure 6, which presents the confusion matrix of predicted versus actual values. Here we observe perfect predictions for males and only one false positive for females.

**Table 3:** Optimal hyperparameters for XGBC and KNC methods.

| Sex | Model | Hyperparameter | Tuned value |
| --- | --- | --- | --- |
| M | XGBC | colsample_bytree | 0.7 |
|  |  | gamma | 0.5 |
|  |  | subsample | 0.7 |
|  |  | learning_rate | 0.1 |
|  |  | max_depth | 3 |
|  |  | n_estimators | 100 |
| F | KNC | metric | manhattan |
|  |  | weights | distance |
|  |  | n_neighbors | 7 |

**Table 4:** Test scores of the classifiers across the considered metrics for males (M) and females (F).

| Accuracy |  | Precision |  | Recall |  | F1 score |  |
| --- | --- | --- | --- | --- | --- | --- | --- |
| M | F | M | F | M | F | M | F |
| 1.00 | 0.99 | 1.00 | 0.94 | 1.00 | 1.00 | 1.00 | 0.97 |

**Figure 6:**
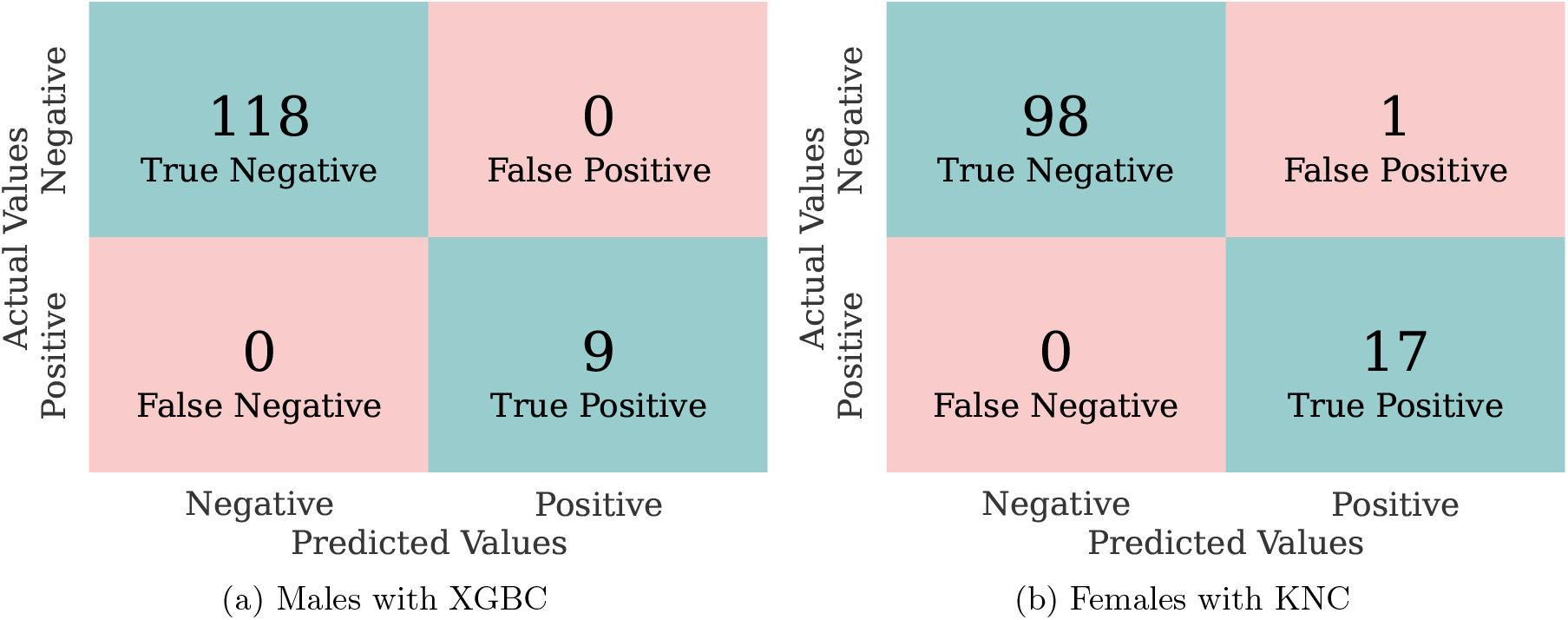
Confusion matrix of the classifiers. True positives (negatives) represent the number of correctly predicted positive (negative) cases. False positives (negatives) indicate the number of incorrect positive (negative) predictions.

##### Gaussian process regression model

The second step of the emulators is based on a GPR model that predicts the ΔQT values. We split our data as done for the classifier and we exclude inputs that are identified as positive. Similarly to the preliminary version of the emulators, we first tune the optimal hyperparameters through an exhaustive search on the training set. These hyperparameters are selected to maximize the R^2^ score of predictions and are reported in Table 5.

**Table 5:** Optimal hyperparameters for the GPR model, for males (M) and females (F).

| Model | Hyperparameters | Tuned value |  |
| --- | --- | --- | --- |
|  |  | M | F |
| GPR | kernel | Matérn 1.5 | Matérn 2.5 |
| | gamma | $10^{-3}$ | $10^{-3}$ |
|  | subsample | 20 | 5 |

The tuned GPR model is used to predict the ΔQT values corresponding to the ionic channel blockades of the test set. Figure 7 shows that the ΔQT predictions are highly accurate both for males and females, a result that is also confirmed in Table 6, where we report the R^2^ score and the prediction errors.

**Table 6:** R^2^ scores and prediction errors on the test set for males (M) and females (F)

| $R^2$ score | | $\epsilon^{\text{MAE}}$ [ms] | | $\epsilon^{\text{MAPE}}$ [%] | | $\epsilon^{\text{RMSE}}$ [ms] | |
| --- | --- | --- | --- | --- | --- | --- | --- |
| M | F | M | F | M | F | M | F |
| 0.998 | 0.996 | 1.491 | 2.107 | 3.063 | 4.433 | 2.582 | 3.083 |

**Figure 7:**
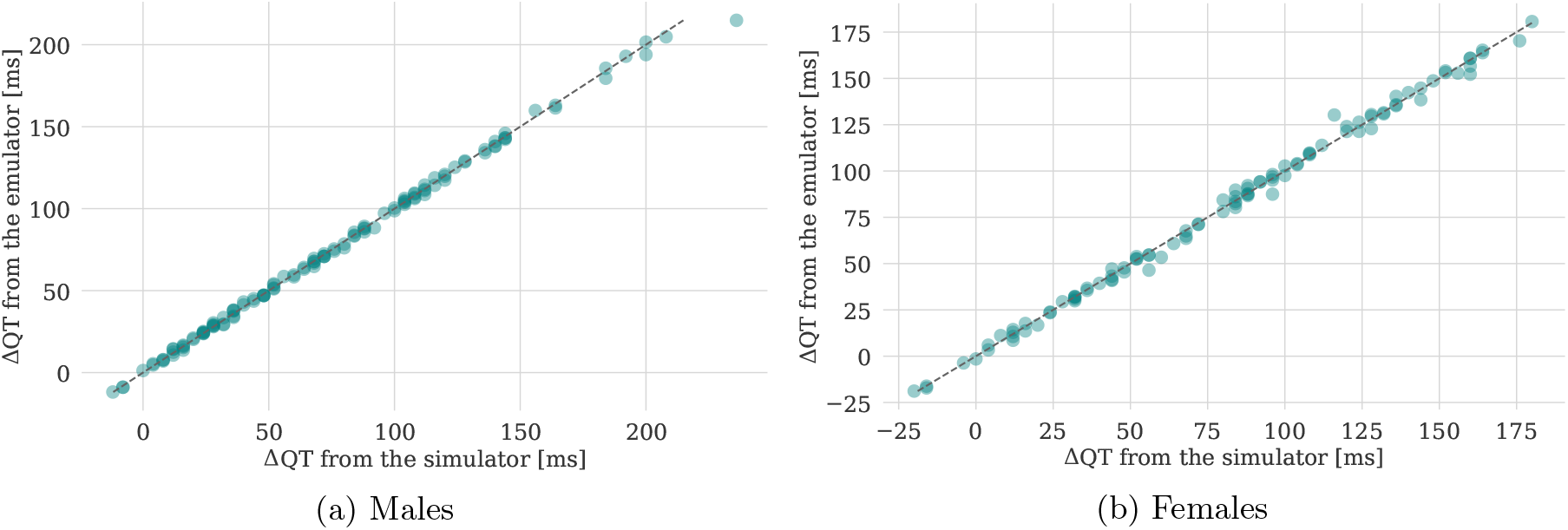
Comparison of ΔQT obtained with the simulator and emulators.

### 2.4 Global sensitivity analysis

After fine-tuning our emulators, we perform again the GSA to address two key questions:

- Which ionic channels mainly influence the output?
- Which ionic channels primarily contribute to a QT prolongation that is classified as high-risk?

The first question is directly determined by the emulator output, while the second is assessed based on whether the predicted ΔQT exceeds the high-risk threshold of 50 ms [47].

Figure 8 shows the GSA results, separated by sex, with the first analysis displayed on the top and the second one on the bottom. In each plot, the solid bars represent the *S*_1_ indices, indicating the main individual effects, while the striped bars indicate the *S*_*T*_ indices, reflecting the total effects, i.e., describing the interactions between different ion channel blockades. Confidence intervals are also included to show the uncertainty in the estimation of the Sobol’ indices. From the top plots, we observe that *I*_Kr_ is the most relevant channel in predicting QT prolongation, regardless of sex. It is important to note that *I*_Kr_ refers to the rapidly activating delayed rectifier potassium current, which is largely mediated by the hERG (human Ether-à-go-go Related Gene) channel, often referred to simply as *I*_Kr_ channel. The other channels have significantly less influence on the output, and interactions between them are negligible since *S*_1_ and *S*_*T*_ are very similar for each channel considered. On the other hand, the second analysis highlights that *I*_Kr_, *I*_NaL_, and *I*_Ks_ are the three most influential channels in contributing to a high-risk QT prolongation. Notably, interactions between different channel blockades are significant, particularly for the most influential input, *I*_Kr_.

**Figure 8:**
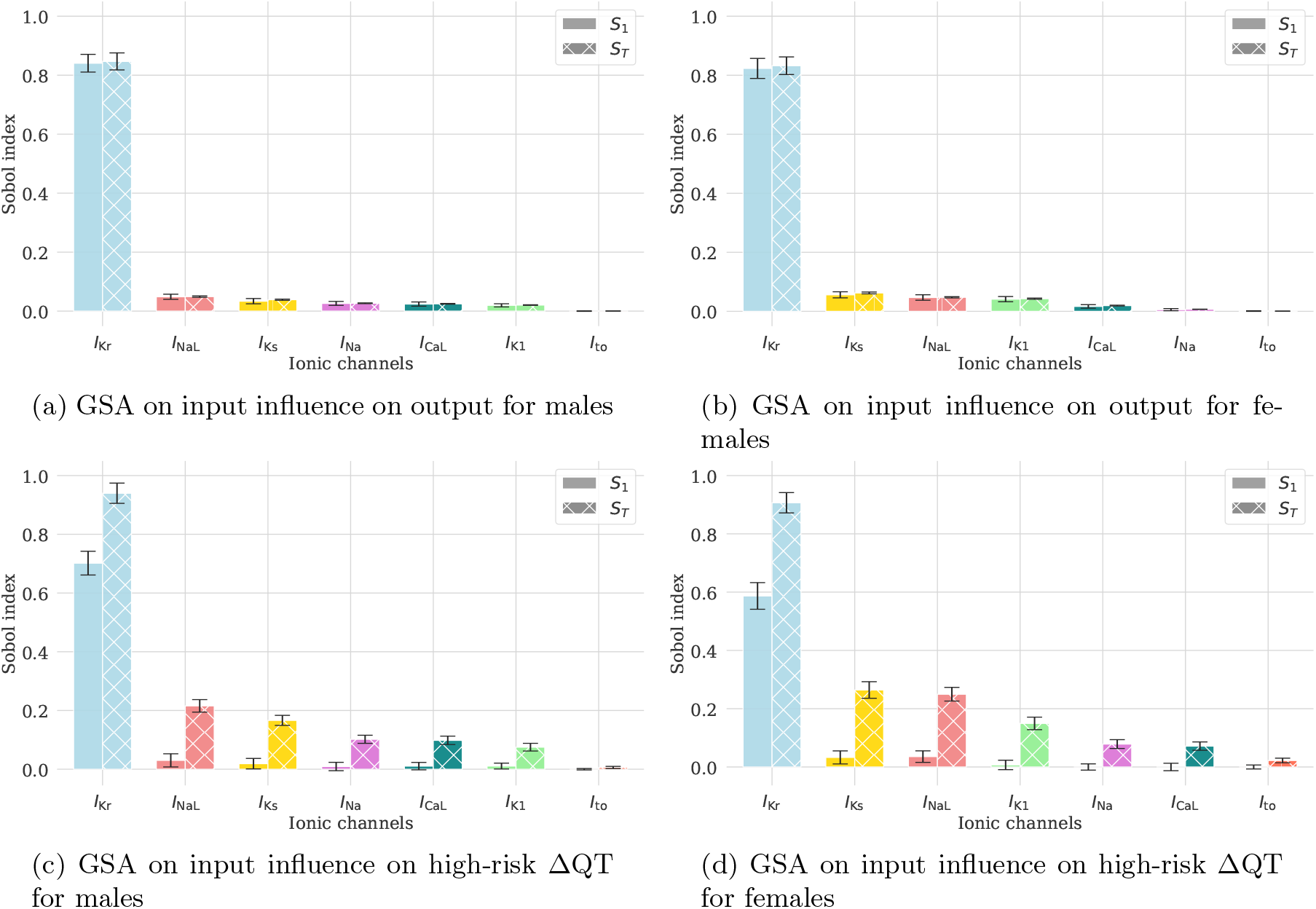
First and total Sobol’ indices highlighting the influence of ionic channel blockades on the predicted ΔQT (top) or in determining if the QT prolongation is of high-risk level (bottom).

### 2.5 Reproducing benchmark drugs with the emulators

In Figure 9, we present the concentration-QT prolongation (C-ΔQT) predictions of the emulators, simulator and clinical data under the influence of dofetilide. The slope of the C-ΔQT relationship is crucial, as it measures the rate of change in the QT interval per unit increase in drug concentration, allowing for comparison of drug effects across different populations.

**Figure 9:**
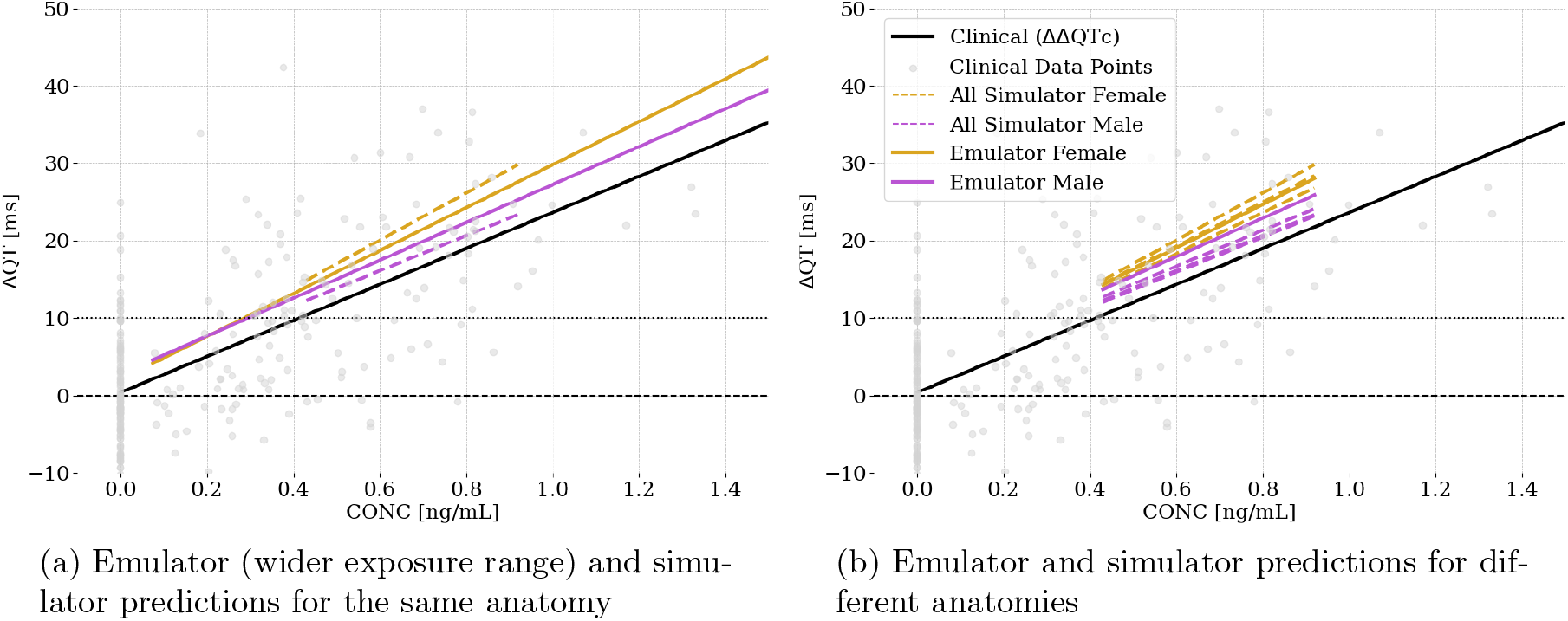
Application of the emulators to predict C-ΔQT response for dofetilide. ΔΔQTc clinical, ΔQT simulated and ΔQT emulated as a function of plasma concentration. The grey dots denote the observed ΔΔQTc with respect to plasma concentration for each measurement taken in the clinical trial extracted from Darpo *et al*. (2015) [53].

Our emulators are developed using single average geometries that represent typical adult anatomies. Consequently, the simulator results shown in Figure 9(a) are based on the same anatomies used for training the emulators. Leveraging the performance of the emulators, we extend the concentration range beyond the typical experimental limits. This illustrates their potential to predict drug effects at higher concentrations in immediate response time, as we will better discuss in Section 2.6. Results demonstrate a strong similarity between the outputs of the emulators and the simulator, confirming the accuracy of the emulators. We calculate the slopes of the C-ΔQT relationships for clinical, simulator, and emulator data, and then compute the relative errors between these slopes as percentages. For females, the relative error between the emulator and simulator slopes is 9.0%, the error between the simulator and clinical slopes is 31.1%, and the error between the emulator and clinical slopes is 19.2%. For males, the relative error between the emulator and simulator slopes is 7.7%, the error between the simulator and clinical slopes is 2.4%, and the error between the emulator and clinical slopes is 5.2%. These results underscore the consistent alignment among all datasets, especially considering the limited female representation in the clinical trial [53]. Then, to assess the robustness of our emulators, we compare their predictions against simulator results from six different anatomical variations, see Figure 9(b). Despite being trained on a single average anatomy, the emulator predictions remain highly accurate across different anatomies. Specifically, for female anatomies, the emulator predictions show relative errors ranging from 1.8% to 8.0% compared to the simulator. For male anatomies, relative errors range between 7.2% and 9.6%. When comparing the emulator slopes to clinical data, we observe relative errors of approximately 20.6% for females and 7.0% for males. The larger error in females is closely tied to the fact that a really small number of women were included in the clinical trial. These results suggest that, even when trained on a single average geometry, the emulators provide predictions that are comparable to the simulator alignment with the usually sparse clinical data. This suggests that the emulators are effective and capable of generalizing well across various anatomical conditions, offering valuable insights despite the inherent variability in clinical data.

To further evaluate the accuracy of the emulators, we predict ΔQT for four benchmark drugs: moxifloxacin, ondansetron, dofetilide, and verapamil. We compare the emulator results with clinical data and with simulator results from our previous study [23], which used six different anatomies. To ensure consistency and avoid bias, we use the same concentration protocol for each drug, involving three ascending concentrations, except for verapamil, which is tested at four concentrations. The calculated C-ΔQT relationships are shown in Figure 10. Our analysis shows that the slopes of the emulator regression lines are remarkably similar to those produced by the simulator across all benchmark drugs. This similarity is observed alongside the expected differences among sexes. Table 7 provides a comprehensive summary of the regression analysis results for each drug, comparing clinical data with the simulator and emulator results. The latter closely mirror the simulator results for all drugs. Specifically, the slopes of the emulator predictions fall within the confidence intervals of the simulated slopes, achieving critical concentrations within a 0.25-fold range of simulated results. This alignment extends to most clinical trends as well, supporting the emulator reliability and accuracy in predicting QT interval prolongation under the effect of different drugs. Ondansetron and moxifloxacin present some discrepancies with respect to the clinical data in slope and intercept, respectively, leading to differences in critical concentrations. Drug experimental variability and clinical data intrinsic biases may justify the observed differences, as also discussed in [23].

**Table 7:** Summary of regression analysis results for four benchmark drugs. For each drug, we present the following metrics: Slope, which quantifies the rate of change in QT prolongation with respect to drug concentration; Intercept, the baseline QT prolongation when drug concentration is zero; Std Err, the standard error of the slope, indicating the precision of the regression estimate; Confidence Interval, providing the range within which the true slope is expected to fall with 95% confidence; and Critical Conc., the drug concentration at which the QT prolongation reaches 10 ms. Results are provided for both clinical data and predictions from the simulator and emulators, differentiated by sex, when applicable.

| Drug | Data Type | Slope | Intercept | Std Err | Confidence Interval | Critical Conc. [ng/mL] |
| --- | --- | --- | --- | --- | --- | --- |
| Dofetilide | Clinical | 23.252 | 0.387 | 1.653 | $\pm 3.239$ | 0.413 |
| | Simulator Female | 24.720 | 1.131 | 0.886 | $\pm 1.737$ | 0.359 |
| | Simulator Male | 24.925 | 0.201 | 0.979 | $\pm 1.918$ | 0.393 |
| | Emulator Female | 28.983 | 1.254 | 0.768 | $\pm 1.505$ | 0.302 |
| | Emulator Male | 26.185 | 1.650 | 1.059 | $\pm 2.076$ | 0.319 |
| Moxifloxacin | Clinical | 0.007 | 1.094 | 0.000 | $\pm 0.001$ | 1356.450 |
| | Simulator Female | 0.007 | 7.125 | 0.000 | $\pm 0.001$ | 427.089 |
| | Simulator Male | 0.008 | 4.625 | 0.001 | $\pm 0.001$ | 699.382 |
| | Emulator Female | 0.009 | 7.216 | 0.001 | $\pm 0.001$ | 325.114 |
| | Emulator Male | 0.008 | 5.766 | 0.001 | $\pm 0.001$ | 546.715 |
| Ondansetron | Clinical | 0.035 | -0.033 | 0.004 | $\pm 0.007$ | 290.285 |
| | Simulator Female | 0.080 | -0.489 | 0.003 | $\pm 0.005$ | 130.858 |
| | Simulator Male | 0.077 | -0.560 | 0.003 | $\pm 0.006$ | 136.854 |
| | Emulator Female | 0.095 | 0.090 | 0.000 | $\pm 0.001$ | 103.977 |
| | Emulator Male | 0.083 | 0.455 | 0.002 | $\pm 0.003$ | 114.323 |
| Verapamil | Clinical | 0.035 | 1.863 | 0.003 | $\pm 0.006$ | 232.144 |
| | Simulator Female | 0.044 | -0.150 | 0.002 | $\pm 0.005$ | 231.871 |
| | Simulator Male | 0.046 | -1.810 | 0.002 | $\pm 0.003$ | 259.001 |
| | Emulator Female | 0.045 | 0.009 | 0.001 | $\pm 0.003$ | 223.479 |
| | Emulator Male | 0.043 | 0.684 | 0.001 | $\pm 0.001$ | 216.686 |

**Figure 10:**
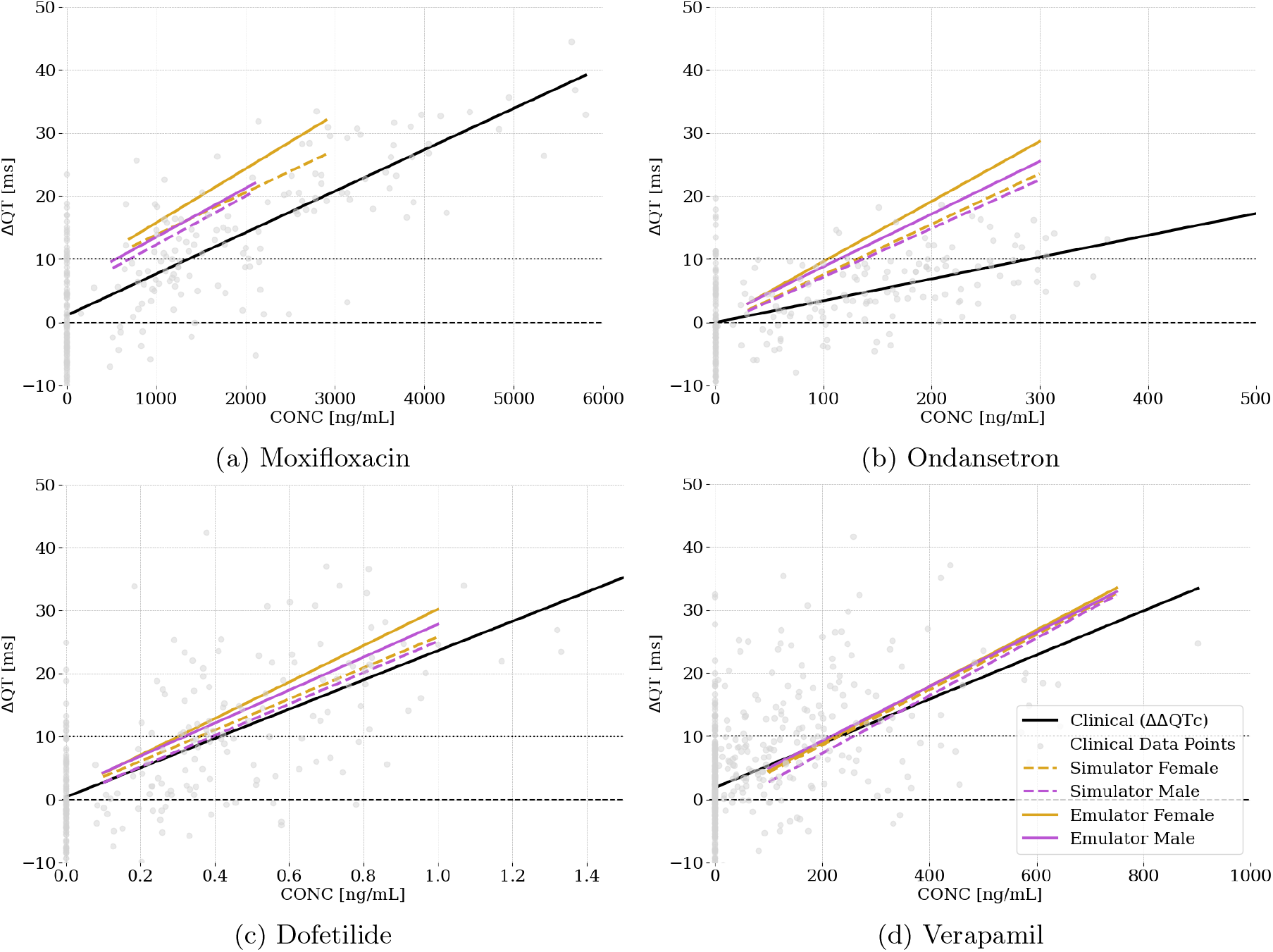
ΔΔQTc clinical, ΔQT simulated and ΔQT emulated as a function of plasma concentration for each drug assessed: moxifloxacin, ondansetron, dofetilide, verapamil. The grey dots denote the observed ΔΔQTc with respect to plasma concentration for each measurement taken in the clinical trial extracted from Darpo *et al*. (2015) [53] and Vicente *et al*. (2019) [54].

### 2.6 Computational performance

We present the results in terms of the computational performance of both the simulator and the emulators. Table 8 shows the computational times required to build the emulators: running 900 simulations took approximately 123 days of CPU time. The time to find the optimal hyperparameters was about 48 minutes, while the emulator training took only seconds. It is important to note that these operations are performed only once.

**Table 8:** Computational times to build the emulators: CPU time to run 900 simulations, time to find the optimal hyperparameters of the emulators, and time to train the emulators. Male and female results are summed. The simulator time has been computed excluding the cases where the initialization of the cellular model did not converge to a periodic solution.

| Simulator<br>(900 cases) | Emulator optimal<br>hyperparameters | Emulator<br>training |
| --- | --- | --- |
| 123 d | 48 min | 23 s |

In Table 9, we present the computational costs associated with reproducing a single case and conducting clinical trials with benchmark drugs using both the simulator and the emulators. A single simulation took, on average, 3.43 h, whereas the emulators can predict the ΔQT in mere centiseconds, providing a speed-up of five orders of magnitude. Generating a clinical trial for a single drug required approximately 9 days of CPU time using the simulator. In contrast, the emulators significantly reduce this cost, delivering accurate results in less than a second.

**Table 9:** Comparison of the simulator and emulator computational performance to predict ΔQT for a single case and to reproduce *in silico* clinical trials of benchmark drugs (average cost for the clinical trial of a single drug). The speed-up is computed as the ratio between reported simulator and emulator times.

|  |  | Simulator | Emulators | Speed-up |
| --- | --- | --- | --- | --- |
| Single case | | 3.43 h | $8 \cdot 10^{-2}$ s | $1.5 \cdot 10^5$ |
| Benchmark drugs | Moxifloxacin | 9 d | $3.2 \cdot 10^{-1}$ s | $2.4 \cdot 10^6$ |
|  | Dofetilide |  |  |  |
|  | Ondansetron |  |  |  |
|  | Verapamil |  |  |  |

## 3 Discussion

We introduced sex-specific emulators capable of predicting drug-induced proarrhythmic risk. Developed separately for each sex, these emulators leveraged a comprehensive dataset derived from high-fidelity 3D electrophysiological simulations, capturing crucial anatomical and phenotypical differences. By taking ionic channel blockades as input, they provided real-time predictions of QT prolongation, ensuring accurate and sex-specific assessments.

To construct our dataset, we leveraged the computational power of Alya [39, 55] to perform 900 electrophysiological simulations (450 per sex), sampling the space of seven ionic channel blockades as input, which totaled approximately 2.1 million CPU hours. This comprehensive dataset, entirely composed of high-fidelity 3D simulations, was essential for developing accurate emulators. Notably, our dataset is comparable in size to that built by Costabal *et al*. [36], who used a mix of 400 low-fidelity (1D) and 45 high-fidelity (3D) simulations.

The usage of 3D cardiac models introduces additional complexity and computational cost compared to 0D single-cell models. However, our analysis demonstrated the clear advantages of 3D modeling for predicting drug-induced proarrhythmic risk, underscoring its importance despite the higher computational demands. 3D models provide greater physiological relevance by capturing spatial heterogeneity and anatomical details crucial for simulating drug effects on cardiac electrophysiology. These models can compute ΔQT, which makes them comparable to clinical trial outcomes, thus enhancing the reliability of computational models for cardiac safety assessment. In contrast, 0D modeling relies on *in vitro* metrics such as APD or qNet, which are not directly translatable to clinical settings. Another significant advantage of 3D models is their capacity to incorporate disease states and comorbidities, particularly through detailed anatomical representations. This is especially valuable because such patients are often excluded from real clinical trials. By using cardiac models, we can ensure that these underrepresented groups are considered in the assessment of drug safety. Our comprehensive dataset allowed us to systematically compare 3D versus 0D modeling in drug proarrhythmic risk assessment. By comparing ΔQT and ΔAPD, we found that while 0D models can generally predict outcomes accurately, their ability to identify electrical propagation abnormalities decreased significantly as the risk level increased. This limitation is concerning because it indicates that 0D models may not reliably predict high-risk scenarios, potentially underestimating arrhythmogenic risks. Furthermore, our comparison between ΔQT and qNet revealed that the latter frequently misclassifies female cell types into incorrect risk categories. This limitation stems from qNet being derived from the original ORd model, which is based on a generalized, endocardial model [41]. As a result, qNet is more accurate for male endocardial cells but fails to account for the sex-specific differences in cardiac electrophysiology that are crucial for accurately assessing risk in females. Additionally, comparing action potentials (from the 0D model) with the ECGs (from the 3D simulation) revealed that similar patterns at the 0D level can lead to different outcomes at the 3D level, especially within the arrhythmogenic window, again emphasizing the importance of 3D models in proarrhythmic risk assessment.

Our study underscored the critical importance of incorporating sex differences in computational models, aligning with previous research findings [21–24]. Our results showed that females are more susceptible to arrhythmias, consistent with earlier studies [25, 56]. The use of 3D electrophysiological models enabled a comprehensive approach to account for sex-specific anatomies and phenotypes. Thus, although 3D cardiac models demand greater computational resources compared to 0D models, their capacity to capture complex physiological details and sex differences – thereby providing more accurate and personalized risk assessments – justifies their use in predicting drug-induced proarrhythmic risk. Thus, our findings emphasized the necessity of high-fidelity 3D simulations to enhance the reliability and accuracy of computational cardiac safety evaluations. One way to mitigate the computational expense of 3D simulations is by developing emulators, or surrogate models, which can predict the outcomes of computationally costly 3D simulations at a fraction of the computational cost.

To compute QT prolongation induced by drugs in real-time, we developed two emulators that utilized a two-step approach: a classifier followed by a GPR model. The classifier is used to filter cases, improving the robustness and accuracy of our emulators by addressing the challenges inherent in cardiac electrophysiological models, which often display bifurcations and discontinuous responses. Ghosh *et al*. [57] noted that such discontinuities could complicate predictions when using GPR models, as they assume smooth and continuous responses to changes in parameters. By implementing the classifier, we effectively segregated non-arrhythmic from arrhythmic cases, thereby minimizing the complexity and discontinuities presented to the GPR model. Additionally, sex-specific ΔQT thresholds were crucial to avoid the introduction of physiological anomalies that could mislead the training process and predictions. This consideration is particularly relevant for Thorough QT studies, where identifying significant QT prolongation is essential for further clinical development. Literature highlights that a QT prolongation exceeding 20 ms is often significant enough to warrant extra monitoring and labeling [58]. Furthermore, patients experiencing TdP usually show more pronounced QT prolongation before its onset. Sex differences add another layer of complexity, with women being more sensitive to QT prolongation [25]. Expert guidelines suggest that QT intervals longer than 550 ms are associated with a high arrhythmic risk [40]. Consequently, our chosen thresholds – 240 ms for males and 196 ms for females – are set conservatively to account for cases that could lead to a QT interval of 600 ms, ensuring that rare but critical extreme cases are also identified. This approach allowed the regressor to work with a more homogeneous and relevant dataset, thereby enhancing the accuracy and reliability of our predictions.

The sex-based emulators demonstrated high accuracy when compared to simulator results, with errors in the range of 2.6 ms, a relatively minor discrepancy given the magnitude of drug-induced changes. These emulators enabled us to replicate clinical trials for four benchmark drugs and validate our findings against *in vivo* data. This consistency underscored the emulator reliability in forecasting drug-induced QT prolongation across varying concentrations and compounds. The emulators effectively captured the sex-based differences in drug response observed with the simulator, which is crucial for developing sex-specific therapeutic strategies, especially when it comes to high QT-prolonging drugs. Notably, emulators facilitated an in-depth exploration of dofetilide responses at higher concentrations without additional computational expense. Standard *in silico* clinical trials typically require around 9 days of CPU time, but the emulators reduced this time to mere fractions of a second, achieving a speed-up of six orders of magnitude in predicting QT prolongation. To assess the robustness of our model across different anatomical contexts, we compared emulator predictions – trained on fixed anatomy – with simulator results using different anatomical models. In addition, different from the emulators, the simulator results are obtained by combining phenotypical variability [59]. The emulators produced results with errors ranging from 1.8% to 9.6%, a level of error we considered acceptable in comparison to the existent interstudy variability in clinical practice [60]. This demonstrated that our emulators maintain robustness and reliability, even when tested against data with a larger variability.

In addition, these emulators are also tools to identify and prioritize influential inputs by carrying out GSA. This analysis was used to achieve two main objectives: first, to study the influence of each input on the ΔQT response; and second, to identify the key inputs that determine whether there is a high risk of QT prolongation. We found that *I*_Kr_ is the primary predictor of ΔQT, consistent with the findings reported in [36]. The results also demonstrate that the seven inputs can be linearly combined to reconstruct the QT prolongation. However, when evaluating clinically relevant quantities, such as assessing whether the inputs lead to a QT prolongation that is classified as high-risk, requires accounting for high-order interactions between the channels, with *I*_Kr_, *I*_NaL_, and *I*_Ks_ being the most significant contributors. These findings underscore the importance of considering multichannel interactions in proarrhythmic risk evaluation, suggesting that a sole reliance on hERG *in vitro* assessment may be insufficient.

It is important to highlight that the high accuracy of our emulators is largely attributable to the quality and reliability of the underlying 3D cardiac simulator [39]. This simulator is the result of nearly two decades of continuous development, research, and an extensive validation process across multiple studies [61–67]. Its development has involved close collaboration with clinicians and pharmaceutical experts, ensuring that it reflects the complexity of human cardiac physiology and drug interactions [23, 68]. Achieving such precise emulator performance would not have been possible without this foundation of well-established computational models, which have been refined through years of iterative improvements and cross-disciplinary expertise [39, 55, 69–71]. As such, while the emulators significantly reduce computational costs, their effectiveness is intrinsically linked to the robustness and accuracy of the simulator from which they are derived.

This study has some limitations. Firstly, we relied solely on ΔQT to estimate proarrhythmic risk, whereas other electrocardiogram features, such as the Tpeak-Tend interval [72], might provide additional insights into drug effects. Secondly, our dataset does not account for variability in phenotypic expression or anatomical differences, which could be crucial for a more comprehensive risk assessment. Future research should focus on incorporating these variabilities to better represent the spectrum of patient responses and further improve the accuracy of our results when compared to simulator results. Additionally, our current models use a fixed heart rate and a uniform activation protocol across all cases. Moreover, clinical trials include placebo groups to eliminate confounding effects that allow them to compute placebo-corrected ΔQT. Further developments consist of including placebo corrections within the computational models. Future directions also involve performing uncertainty quantification, especially on parameters such as *IC*50, *h* and the free fraction of the drug. Incorporating variability in these parameters could help refine uncertainty ranges in model predictions and enhance the overall robustness of the emulators. An interesting application of the emulators would be to integrate ionic channel margin distributions into QT prolongation risk assessments, as suggested by Leishman *et al*. [73]. This approach would enable the definition of safety margins by accounting for the variability observed in *in vitro* assay-measured parameters. By doing so, it could serve as an effective filter in early drug development, providing a robust sensitivity analysis that informs decision-making processes and better characterizes the risk landscape.

To conclude, our emulators provide a valuable tool for evaluating QT prolongation from the discovery phase and the very early stages of drug development, offering high accuracy in simulating drug-induced effects on cardiac physiology. Serving as an instant-response preliminary design tool for computational clinical trials, these emulators create a virtual trial environment where outcomes can be predicted and concentration ranges refined – all before committing to computationally expensive full-scale simulations. However, it is crucial to underscore that the emulators do not replace the simulator. The simulator remains indispensable, especially in later stages of development, due to its ability to account for a wider array of physiological factors that the emulators, particularly given their current limitations, cannot fully capture. This distinction is vital, as the emulators are primarily intended to complement the simulator by accelerating early-stage assessments, not to substitute the comprehensive analyses that only the simulator can provide. Moreover, the use of surrogate models allows for the comprehensive execution of studies with uncertainty quantification across all variable drug-related parameters, enabling thorough compound evaluation, informed early-stage decision-making, and the efficient design of 3D virtual trials with a larger control over experimental conditions. Furthermore, emulators also address the growing concern of climate impact linked to preclinical and clinical studies, an item that is getting higher on the agenda of large pharmaceutical companies. The emulators unleash the potential to analyze, investigate, and innovate at a marginal/insignificant cost for the environment, representing a negligible carbon footprint. By integrating high-fidelity simulations with immediate prediction capabilities, our emulators advance the concept of digital twins, driving precision medicine forward and improving the efficiency and sustainability of drug development.

## 4 Methods

### 4.1 The electrophysiological simulator

#### Anatomical models

We collect retrospective data from the Visible Heart Lab library at the University of Minnesota [74]. Specifically, we consider two biventricular cardiac geometries (male and female) reconstructed from high-resolution MRI scans. We use human heart anatomies from adult deceased donors with no history of cardiac disease and anatomically normal ventricles in order to represent average healthy individuals.

#### The electrophysiological model

To model cardiac electrophysiology, we consider the monodomain model [75] coupled with the ORd cellular model [41]. We selected the ORd model because it is recognized as the consensus *in silico* model by the CiPA initiative [76]. Specifically, we used a modified version of the ORd model by Passini *et al*. [77] with modified conductances as described in Dutta *et al*. [76]. The location of the activation points was instead set following the work by Durrer *et al*. [78], as we explained in our previous study [63]. Cardiac fibers are modeled using the outflow tract rule-based method [79]. This method is particularly suited for our study due to its ability to accurately assign fiber directions within high-resolution biventricular geometries, which include complex structures such as trabeculae and papillary muscles [63, 79]. We incorporate transmural myocyte heterogeneity by assigning distinct electrophysiological and cellular properties to various regions of the myocardium: endocardial (inner, 30%), mid-myocardial (middle, 40%), and epicardial (outer, 30%) cells. A larger diffusion is assigned to a one-element layer on the endocardial surface to account for the fast conduction of the Purkinje fibers. We set a constant heart rate of 60 bpm.

To generate male and female phenotypes, we apply sex-specific ion channel subunit expression as described in [23, 80, 81] to the two sex-specific anatomies considered in the study.

#### Modeling drugs

To model the effect of drugs, we use a multi-channel conductance-block formulation [9, 23]. Given the conductance *g*_k_ of one of the seven most influent ionic channels *I*_CaL_, *I*_NaL_, *I*_to_, *I*_Ks_, *I*_K1_, *I*_Na_, and *I*_Kr_ [38], we define the ion channel conductance after the drug administration with the following Hill model [9]:

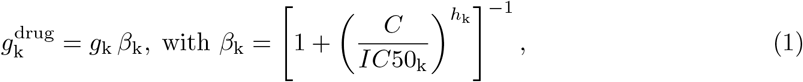

with k = CaL, NaL, to, Ks, K1, Na, Kr. In the equations above, 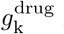 is the conductance of the k–th channel after drug administration, *C* the drug concentration, *IC*50_k_ the concentration required to have a 50% current blockade of the k–th channel, and *h*_k_ the corresponding Hill exponent. Thus, the blockade *β*_k_ of channel k is identified by the Hill parameters *h*_k_, the *IC*50_k_ and the drug concentration *C*. Notice that *C* is defined as the concentration of the drug in the plasma that is not bound to plasma proteins and is therefore available to exert a pharmacological effect. Therefore, it is expressed as

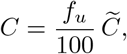

where 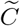 is the total concentration of the drug in the plasma and *f*_*u*_ – expressed in percentage – is the free fraction of the drug: the ratio of the unbound drug concentration to the total drug concentration.

#### Initialization of the electrophysiological model

The 3D simulation is initialized by solving the 0D ORd model for each cell type until the intracellular calcium concentration converges to a periodic solution. Periodicity is determined when the RMSE between consecutive beats is smaller than 10^−7^ μmol for three consecutive beats. The results from the 0D model then serve as initial conditions for the 3D simulation. Cases where periodicity is never achieved are excluded from our analysis (0 cases for males, 37 for females).

For the analysis in Section 2.2, we compared the output from the initialization of the 0D ORd model against the ΔQT obtained in the subsequent 3D simulation.

#### Pseudo ECG and ΔQT computation

To compute the ECG, we assume isotropic electrical conductivity in the torso and we rely on a pseudo-ECG approach, for which the potential in a given generic point ***x***_*_ of the body (where the electrode is positioned) is computed as [82]:

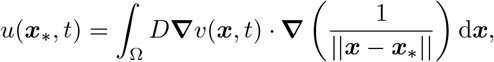

where Ω × (0, *T*) is the spatio-temporal domain, Ω represents the biventricular geometry and *T* is the simulation final time. The function *v* : Ω × (0, *T*) → ℝ is the transmembrane potential and *D* is the orthotropic tensor of local diffusivities. Three leads are then computed as [63]:

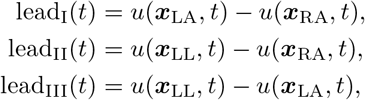

where ***x***_LA_, ***x***_RA_, and ***x***_LL_ are the positions of the electrodes at the left arm (LA), right arm (RA), and left leg (LL), respectively. We refer the interested reader to [63] for additional information on how we compute the pseudo-ECG. The latter is used to compute the QT interval duration at a baseline configuration (QT^bsl^) and after drug administration (QT^drug^). This computation is done automatically for all the cases using an automatic algorithm as we explain in [23]. The QT prolongation is then defined as

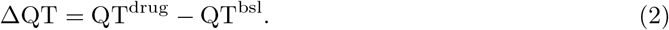

Our computational approach isolates and quantifies the direct impact of drugs on ΔQT, eliminating the confounding effects present in clinical trial cohorts, such as varying patient demographics, concomitant medications, and individual physiological differences. Consequently, our computational data does not require placebo correction, as it inherently excludes these variables. This approach ensures that the observed ΔQT effects are solely due to the drug itself, providing a clearer comparison to clinical data where placebo adjustments account for such confounders [43]. In addition, since here we use a heartbeat period of 1 s (60 bpm), the ΔQT that we compute (from both simulator and emulators) does not require heart rate correction as there is no heart rate variability to account for. This means that the ΔΔQTc from clinical data is directly comparable to the ΔQT from the simulator and emulators.

#### Computational aspects

The computational model is implemented in the multiphysics and multiscale finite element library Alya [39, 55], developed at the Barcelona Supercomputing Center and ELEM Biotech SL. Alya is optimized for efficient execution on supercomputers within a high-performance computing framework. Our study uses biventricular meshes comprising approximately 58 million tetrahedral linear elements, with an average mesh size of 300 μm, and a constant time-step size of 20 μs. Simulations are conducted on the Nord3 machine at the Barcelona Supercomputing Center, simulating three beats with a period of 1 s each. The computational cost of a single simulation, utilizing 672 cores, is approximately 3.5 h. To develop our emulators, we perform 450 simulations per sex, totaling 900 simulations and corresponding to 2.1 million CPU hours globally.

### 4.2 The emulators

To assess real-time QT prolongation caused by drugs, we develop sex-specific emulators that integrate a classifier followed by a regressor. The classifier enhances the robustness and accuracy of our emulators by filtering out non-arrhythmic and arrhythmic cases, reducing dataset complexity and discontinuity. This is crucial given the bifurcations and discontinuous responses in cardiac electrophysiological models, as noted by Ghosh *et al*. [57]. We also employ sex-specific ΔQT thresholds to avoid physiological unfeasible prolongations that could mislead the training and prediction processes. Our thresholds of 240 ms for males and 196 ms for females, which are designed to account for a QT interval of 600 ms, provide a conservative approach to identifying at-risk individuals. This methodology ensures the regressor operates on a more homogeneous subset of data, enhancing the reliability of the predictions.

Let 𝒟 = (***X, y***) be a dataset of size *N*, where ***X*** = (***x***_1_, *…*, ***x***_*N*_) is the input matrix, and ***y*** = (*y*_1_, *…, y*_*N*_) is the output vector. The input of our model are the current blockages (1 − *β*_k_) defined in Equation (1), with k = CaL, NaL, to, Ks, K1, Na, Kr. The output consists of a binary vector for the classifier (where 1 denotes an arrhythmic ECG and 0 non-arrhythmic ECG) and the ΔQT, computed as in Equation (2), for the regressor. This results in a dataset of the following size: ***X*** ∈ ℝ^7×*N*^, ***y*** ∈ ℝ^*N*^. We begin by splitting the dataset 𝒟 into two parts: the training set 𝒟^train^ and the test set 𝒟^test^. The training set, comprising 72% of the data selected randomly, is used to tune the hyperparameters of both the classifiers and the regressor. The remaining 28% forms the test set, which is used to evaluate their performance. To ensure reproducibility and consistency across the consecutive steps of the emulators, we use the same sets 𝒟^train^ and 𝒟^test^ for classifiers and regressors. The input data are standardized in each step of the procedure, while the output vectors need to be normalized only for the regressor step. Data normalization is performed using the S`tandardScaler` package from the s`cikit-learn` Python library [83], which removes the mean and scales the data to unit variance.

To tune the optimal hyperparameters of the emulators, and to evaluate their performance, we employ a nested *k*-fold cross-validation strategy on the training set [84], with *k* = 5.

#### 4.2.1 Classifier

We considered several classifiers, ranging from Random Forests, XGBC, Linear Regression, Support Vector Machine, and KNC. Of these, XGBC provided the most accurate results for males, while KNC performed best for females. For a detailed explanation of these methodologies, refer to [50, 51].

##### Tuning of classifier hyperparameters and performance analysis

The tuning of the optimal hyperparameters for the XGBC and the KNC methods is performed through the s`cikit-learn` library and the G`ridSearchCV` package. For XGBC and KNC we seek to determine the optimal hyperparameters among the possibilities reported in Table 10.

**Table 10:**
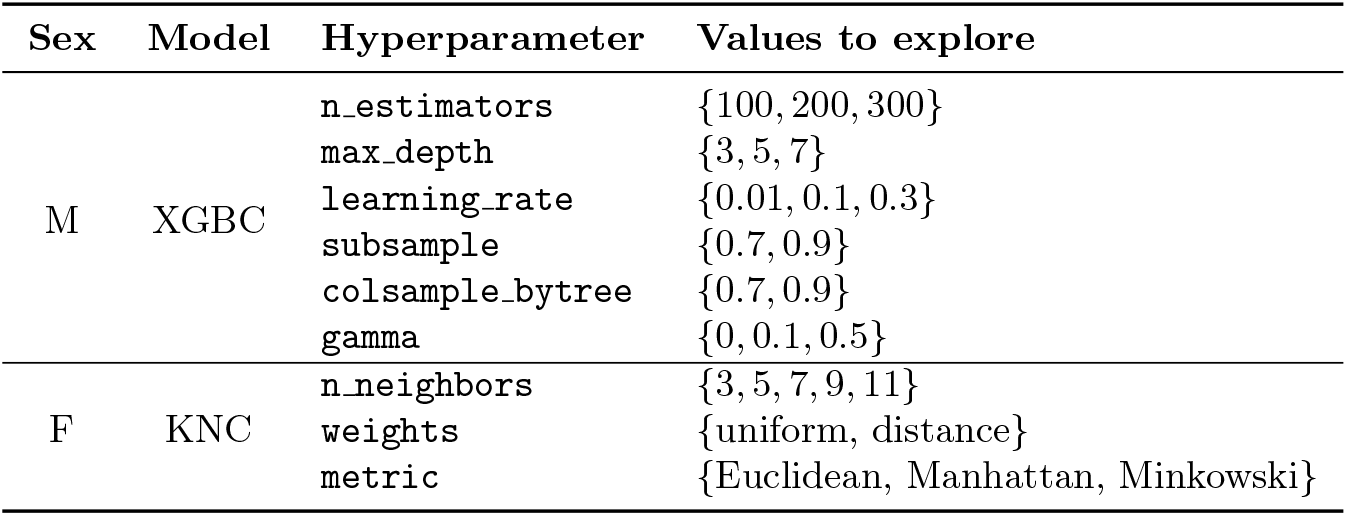
Hyperparameters description and set of selected possible values to explore with the automated exhaustive search for XGBC and KNC methods.

To define the evaluation metrics, we first introduce the following terminology to categorize the classifier outcomes:

*True Positive* (TP): predicted values correctly identified as positive.

*True Negative* (TN): predicted values correctly identified as negative.

*False Positive* (FP): predicted values identified as positive, while the actual ones are negative.

*False Negative* (FN): predicted values identified as negative, while the actual ones are positive.

Table 11 presents the metrics used for evaluation. Specifically, the optimal hyperparameters are selected to maximize the F1 score, as it is the most representative metric for our purposes. Indeed, the F1 score balances the impact of both false negatives and false positives, which are critical factors in medical applications, and offers greater robustness when dealing with imbalanced datasets [85]. After selecting the optimal hyperparameters, we use the test set to evaluate the quality of predictions on unseen data.

**Table 11:** Metrics used to evaluate the classifiers’ performance [52].

| Accuracy | Precision | Recall | F1 score |
| --- | --- | --- | --- |
| $\frac{TP + TN}{\# \text{ predictions}}$ | $\frac{TP}{TP + FP}$ | $\frac{TP}{TP + FN}$ | $\frac{2TP}{2TP + FP + FN}$ |

#### 4.2.2 Gaussian process regression model

Here we focus on the regression model, whose goal is to capture the underlying relation between inputs and outputs. This model is designed to provide efficient approximations of the outputs for novel input data that are not contained in the training set. We evaluated several machine learning methods, including GPR model, multi-layer perceptron regressor, random forests, and XGBoost regression. Among these, the GPR model yielded the most accurate results. Detailed information on GPR models can be found in [86] and their application in cardiac modeling in [27, 29, 31].

##### Tuning of regressor hyperparameters and error analysis

Consider again the training set 𝒟^train^ and the test set 𝒟^test^. GPR model calibration consists of tuning the hyperparameters of the kernel function on 𝒟^train^. This is done by maximizing the log-marginal-likelihood, that is the probability of reproducing the given output values with the emulators.

The tuning of the hyperparameters of the emulators is based again on a nested 5-fold crossvalidation strategy on the training set. We perform the exhaustive search with the G`ridSearchCV` class, which aims to find the optimal hyperparameters among the possibilities shown in Table 12.

**Table 12:** Hyperparameters description and set of selected possible values to perform the automated exhaustive search for GPR model. See [86, Section 4.2] for more details on kernel functions.

| Hyperparameter | Values to explore |
| --- | --- |
| kernel | {RBF, Matérn 1.5, Matérn 2.5} |
| alpha | $\{10^{-5}, 10^{-3}, 10^{-2}, 10^{-1}, 1\}$ |
| n_restarts_optimizer | {0, 5, 10, 20, 50} |

**Table 13.** *IC*50, expressed in nmol*/*L, is the concentration of the drug that inhibits 50% of its target ion channel activity; *h* (Hill coefficient, dimensionless) describes the steepness of the drug’s concentration-response curve. The ionic block profile refers to the sources of these two parameters. 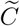 is the total concentration of the drug in the plasma, and *f*_*u*_ is the free fraction of the drug. For Moxifloxacin, different concentrations are used for males and females, according to the reported higher maximum concentrations observed in females [90]. The concentrations were selected based on reported concentrations in clinical trials. Different concentrations are used for the study in Figure 9 and the benchmark study in Figure 10, as the first one is intended at understanding the emulator performance in different physiological conditions, while the latter aims to provide a comprehensive evaluation of drug safety profiles.

|  |  | Moxifloxacin | Ondansetron | Dofetilide | Verapamil |
| --- | --- | --- | --- | --- | --- |
| $I_{CaL}$ | $IC50$ | - | 22551 | - | 202 |
| | $h$ | - | 0.8 | - | 1.1 |
| $I_{Kr}$ | $IC50$ | 93041 | 1492 | 4.9 | 499 |
| | $h$ | 0.6 | 1.0 | 0.9 | 1.1 |
| $I_{K1}$ | $IC50$ | - | - | - | - |
| | $h$ | - | - | - | - |
| $I_{to}$ | $IC50$ | - | - | 18.8 | - |
| | $h$ | - | - | 0.8 | - |
| $I_{Ks}$ | $IC50$ | 50321 | - | - | - |
| | $h$ | 1.0 | - | - | - |
| $I_{NaL}$ | $IC50$ | 382337 | 19181 | - | - |
| | $h$ | 1.1 | 1.0 | - | - |
| $I_{Na}$ | $IC50$ | - | - | - | - |
| | $h$ | - | - | - | - |
| Ionic block profile |  | [38] | [38] | [76] | [38] |
| $\tilde{C}$ [ng/mL] | | 700, 1500, 2900 (F) | 30, 100, 300 | 0.43, 0.92 (Fig. 9) | 99.6, 199.2, 398.4, 750 |
|  |  | 500, 1000, 2100 (M)<br>(Fig. 10) | (Fig. 10) | 0.1, 0.3, 1 (Fig. 10) | (Fig. 10) |
| $f_u$ [%] | | 65 | 27 | 35 | 12 |

To define the metrics used to tune the optimal hyperparameters and evaluate the model performance, we introduce some notation. Given the input data ***X***^test^, we denote by ***y***^pred^ the corresponding prediction of ΔQT intervals. The ground-truth output vector is denoted by ***y***^test^. We select the hyperparameters that maximize the R^2^ score, defined as

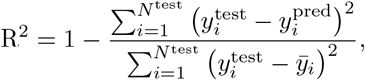

where 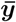 is the average of ***y***^test^. Evaluating the accuracy of the emulators involves comparing the predicted values with the actual ones. To this end, we use the following error metrics, namely the mean absolute error (MAE), the mean absolute percentage error (MAPE), and the root mean squared error (RMSE), defined as:

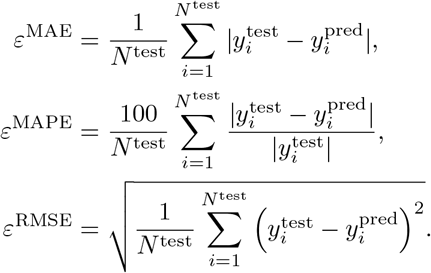

Finally, we evaluate the performance of the model with the optimized hyperparameters on the test set.

### 4.3 Global sensitivity analysis

In this work, we perform GSA [49, 87] twice: firstly to find the most influential channels from the preliminary emulators, allowing us to later enhance the database with higher blockades; secondly, to identify and prioritize influential inputs from our enhanced emulators. GSA is divided into the following steps:

1. *Sampling with Sobol’ sequences* : Sobol’ sequences are quasi-random sequences that ensure a more even and thorough exploration of the parameter space compared to traditional random sampling methods. We use these sequences to generate a set of diverse input parameter combinations (ionic channels’ blockades). Specifically, we sample 2^12^ = 4096 points in the parameter space (this number is typically chosen as a power of 2 to ensure robust coverage and convergence properties).
2. *Model evaluation*: For each sampled parameter combination, we evaluate our emulators to obtain corresponding model outputs (ΔQT).
3. *Variance decomposition and sensitivity indices* : Sobol’ sensitivity analysis decomposes the total variance of the model output into contributions from individual parameters (first-order sensitivity indices, *S*_1_) and their interactions (total-order sensitivity indices, *S*_*T*_). This decomposition provides quantitative measures of the relative influence of each parameter and interaction on the variability of the model output.

We run GSA simulations using the SALib Python library [88, 89], performing two key analyses. The first examines how input parameters influence the emulator outputs (i.e., ΔQT), while the second identifies the most influential factors contributing to QT prolongation risk. In this case, after computing the ΔQT, we categorize the results into a binary vector, labeling them as high or low risk based on whether the predicted ΔQT exceeds the specified high-risk threshold of 50 ms [47].

For the preliminary version of the emulators, we focus on the second analysis. Using Sobol’ sequences, we perform the GSA to identify the three ionic channels that have the most significant impact on the drug-induced high risk of QT prolongation. The primary objective is to isolate regions within the sample space that surround the thresholds for arrhythmic events, thereby enhancing the training dataset in this critical area. Sobol’ sensitivity analysis is particularly well-suited for this purpose due to its variance-based approach, which systematically decomposes the total variance in model output into contributions from individual parameters and their interactions.

After developing and tuning the final version of the emulators, we repeat the GSA to further explore the impact of ionic channel blockades on QT prolongation. The enhanced version of the emulators is used to perform both the analyzes outlined above.

### 4.4 Reproducing benchmark drugs with the emulators

We evaluate the emulator performance by comparing their predictions of C-ΔQT for various benchmark drugs with simulator results and publicly available clinical data [53]. The simulator results, shown in our analysis, follow the population approach outlined in [59], which incorporates phenotypic variability, as we presented in [23]. Additionally, the simulator incorporates anatomical variability using data from the Visible Heart Lab at the University of Minnesota [74], which features a comprehensive collection of patient cases with varying BMI, age, and anatomical characteristics. In contrast, our emulators do not account for any of these sources of variability.

To assess the emulator accuracy, we perform linear regressions on the C-ΔQT relationships derived from clinical data, simulator results, and emulator results. For the clinical data, the linear regression is based on ΔΔQTc. This validation analysis is divided into two parts, corresponding to Figures 10 and 9 respectively. Table 13 details the blockades and concentrations used as input data for both of them.

## Author contributions statement

P.D., A.Z., L.B., and C.B. contributed equally to the conception, design, data acquisition, implementation of the computer code and supporting algorithms, analysis, validation, results visualization, writing of the original manuscript, and its subsequent review. B.D. assisted with study design, clinical validation, and manuscript review. C.M. and M.V. contributed to funding acquisition, provided computational resources, and reviewed the manuscript. J.A. supervised the work and contributed to the manuscript review.

## Data availability statement

The data generated and analyzed during this study are proprietary to ELEM Biotech and cannot be publicly shared due to commercial confidentiality. However, access to the data may be considered on a case-by-case basis. Requests for access can be directed to ELEM Biotech compliance department, and will be subject to a data use agreement ensuring compliance with relevant regulations and restrictions.

